# Bridging the Gap: Integrating Cutting-edge Techniques into Biological Imaging with deepImageJ

**DOI:** 10.1101/2024.01.12.575015

**Authors:** Caterina Fuster-Barceló, Carlos García López de Haro, Estibaliz Gómez-de-Mariscal, Wei Ouyang, Jean-Christophe Olivo-Marin, Daniel Sage, Arrate Muñoz-Barrutia

## Abstract

This manuscript showcases the latest advancements in deepImageJ, a pivotal Fiji/ImageJ plugin for bioimage analysis in the life sciences. The plugin, known for its user-friendly interface, facilitates the application of diverse pre-trained neural networks to custom data. The manuscript demonstrates a number of deepImageJ capabilities, particularly in executing complex pipelines, 3D analysis, and processing large images.

A key development is the integration of the Java Deep Learning Library (JDLL), expanding deepImageJ’s compatibility with various deep learning frameworks, including TensorFlow, PyTorch, and ONNX. This allows for running multiple engines within a single Fiji/ImageJ instance, streamlining complex bioimage analysis tasks.

The manuscript details three case studies to demonstrate these capabilities. The first explores integrated image-to image translation and nuclei segmentation. The second focuses on 3D nuclei segmentation. The third case study deals with large image segmentation.

These studies underscore deepImageJ’s versatility and power in bioimage analysis, emphasizing its role as a critical tool for life scientists and researchers. The advancements in deepImageJ bridge the gap between deep learning model developers and end-users, enabling a more accessible and efficient approach to biological image analysis.

The advancements in deepImageJ, detailed in this paper, represent a significant leap in bioimage analysis, crucial for life sciences. By enhancing this Fiji/ImageJ plugin, the research bridges the gap between complex deep learning models and practical applications, making advanced bioimage analysis accessible to a broader audience. This integration of the Java Deep Learning Library (JDLL) within deepImageJ is particularly noteworthy, as it expands compatibility with leading deep learning frameworks. This allows for the seamless execution of multiple models in a single instance, simplifying the construction of complex image analysis pipelines. The implications of this research are far-reaching, extending beyond academic circles to potentially impact various sectors, including healthcare, pharmaceuticals, and biotechnology. The enhanced capabilities of deepImageJ in handling intricate pipelines, 3D analysis, and large images facilitate detailed and efficient analysis of biological data. Such advancements are vital for accelerating research and development in medical imaging, drug discovery, and understanding complex biological processes. This manuscript contribution to the field of bioimage analysis is significant, offering a tool that empowers researchers, irrespective of their computational expertise, to leverage advanced technologies in their work. The wide applicability and ease of use of deepImageJ have the potential to foster interdisciplinary collaborations, drive innovation, and facilitate discoveries across various scientific and industrial sectors.

## 1. Introduction

Bioimage analysis has undergone remarkable progress due to the advent of open-source tools, making advanced technologies more accessible. These tools include BiaPy^(1)^, CellPose^(2)^, CellProfiler^(3)^, Icy^(4)^, Ilastik^(5)^, ImJoy^(6)^, Napari^(7)^, QuPath^(8)^, ZeroCostDL4Mic^(9)^ and others. One notable development in this field is the emergence of zero-code tools, which streamline the integration of complex analysis pipelines and Deep Learning (DL) networks across various bioimage analysis domains. A key example is Fiji/ImageJ^(10)^, an open-source desktop application central to bioimage analysis. It offers extensive capabilities enhanced by a vibrant community that develops plugins, enabling tasks ranging from basic image processing to advanced DL networks specialized in star-convex object segmentation (i.e., nuclei)^(11)^.

DeepImageJ^(12)^, a freely available plugin for ImageJ, stands out in the realm of zero-code toolkits. It provides an integrated environment within ImageJ for executing third-party models from DL libraries. Notably, deepImageJ is a recognized community partner of the BioImage Model Zoo (bioimage.io)^(13)^, hosting pre-trained models for life sciences. Its user-friendly installation process simplifies complex DL pipeline applications for biologists. DeepImageJ’s practicality is demonstrated in effective workflows for microscopy image analysis^(14)^.

This manuscript presents the latest version of deepImageJ, leveraging the BioImage Model Zoo’s strengths. It introduces the Java-based Deep Learning Library (JDLL), marking a significant advancement in accessible tools for bioimage analysis. The new version, deepImageJ 3.0, brings notable improvements, including enhanced integration with the BioImage Model Zoo and increased compatibility with various DL frameworks. These advancements make deepImageJ a more versatile and powerful tool in the Fiji/ImageJ ecosystem, especially in the application of complex image analysis tasks. The case studies included in this manuscript exemplify the practical applications of these improvements, showcasing deepImageJ 3.0’s enhanced capabilities in diverse bioimage analysis scenarios.

## 2. Advancements in deepImageJ 3.0: Expanding Capabilities in Bioimage Analysis

With the recent update of deepImageJ (deepImageJ 3.0), a range of significant advancements have arisen, thereby expanding deepImageJ’s functionalities and broadening its applicability in bioimage analysis. These new features are designed to simplify the integration and execution of DL models, offering researchers a more versatile and efficient toolset.

### 2.1 Java Deep-Learning Library: A Comprehensive Toolkit

A pivotal feature of deepImageJ 3.0 is its integration with the JDLL^(15)^. JDLL acts as an all-encompassing toolkit and application programming interface, facilitating the creation of sophisticated scientific application and image analysis pipelines with DL functionality. This library simplifies the complex tasks of installing, maintaining, and executing DL models, with support for major frameworks like TensorFlow, PyTorch, and ONNX. The DL engine installer and DL model runner within JDLL provide an intuitive workflow for downloading, integrating and performing inference, offering a harmonized approach to utilizing various DL frameworks. This synergy between deepImageJ and JDLL significantly enhances the ability to execute DL models within Fiji/ImageJ, offering researchers a more streamlined and cohesive environment for bioimage analysis^(15)^.

### 2.2. Multiple Engine Compatibility: Running Different Engines in a Single Fiji/ImageJ Instance

A noteworthy advancement in deepImageJ 3.0 is its newfound capability to load and unload multiple DL frameworks within the same Fiji/ImageJ instance. This development allows for the building of image analysis pipelines that incorporate multiple DL stages, utilizing different engines. This improved compatibility enables seamless integration of models developed in TensorFlow, PyTorch, and ONNX, creating a unified workflow. Such an enhancement provides users with the flexibility to execute a variety of models in a single, integrated pipeline. An example of this is presented in Case Study 1, where image-to-image translation and cell segmentation are performed concurrently within the same Fiji/ImageJ instance.

### 2.3. Extended Framework Compatibility: Supporting Various Versions of DL Frameworks

DeepImageJ 3.0 expands its compatibility with a variety of DL frameworks, including different versions. Now, users can run models created with TensorFlow 2 and ONNX, alongside the already supported TensorFlow 1 and PyTorch. This extension in compatibility broadens the range of executable models accessible within the BioImage Model Zoo ecosystem. Researchers can take advantage of a more diverse array of DL frameworks and their versions, thereby enriching the diversity of models available from bioimage analysis.

### 2.4. Handling Large Images: Leveraging ImgLib2

DeepImageJ 3.0, built upon ImgLib2, is equipped to handle large images, thereby demonstrating its capacity for processing extensive datasets within some limits depending on the computer’s capability. For instance, processing a single large image may require as much as one-tenth of the computer’s RAM capacity. The integration of ImgLib2 significantly boosts the flexibility and scalability of deepImageJ, making it adept at accommodating the large image sizes often encountered in bioimage analysis. This feature together with its tiling strategy^(12)^, ensures that researchers have the ability to apply DL models to a wide array of image data, enabling thorough and detailed analyses.

## 3. Case Studies

In this section, we showcase real-world applications of deepImageJ 3.0 through a series of case studies. These studies highlight the software-enhanced features, as previously discussed. Through these practical examples, our goal is to demonstrate the significant impact and versatility of deepImageJ in tackling various challenges encountered in bioimage analysis.

### 3.1. Case Study 1: deepImageJ Pipeline for Integrated Image-to-Image Translation and Nuclei Segmentation

This case study showcases the advanced capabilities of deepImageJ, particularly its proficiency in integrating and executing diverse DL approaches, often challenged by library and dependency incompatibilities within a typical Python environment. Specifically, we have developed a sophisticated bioimage analysis pipeline that combines the creation of artificially labeled nuclei images from membrane staining images with subsequent nuclei segmentation. This approach allows us to generate synthetic nuclei images, which are easier to process, from an input image (stained cell membranes) that is more suitable for live imaging due to lower phototoxicity compared to direct nuclear staining.

DeepImageJ 3.0 utilizes its enhanced features to combine two distinct DL networks: Pix2Pix^(16)^ and StarDist^(17)^. This integration enables the conversion of membrane-stained images to nuclei stains using Pix2Pix, exported in Pytorch 2.0.1, followed by nuclei segmentation with StarDist, implemented in TensorFlow 2.14. This case study not only demonstrates deepImageJ’s capacity to integrate diverse approaches but also highlights its ability to run models with different engines, addressing the often encountered incompatibility of libraries and dependencies in Python environments^(9)^. The pipeline effectively manages and executes these models, each requiring distinct engines, showcasing deepImageJ’s versatility in handling complex bioimage analysis tasks.

To showcase the versatility of deepImageJ, the initial step employs Pix2Pix, a conditional generative adversarial network designed for image-to-image translation. This network is used to convert fluorescence microscopy images of Lifeact-RFP (actin, membrane stain) to sir-DNA (DAPI, nuclei stain) images. Subsequently, the pipeline applies nuclei segmentation using StarDist, showcasing a comprehensive analysis in a unified workflow.

The construction of this pipeline is depicted in Figure 1. It comprises two main phases (i) training both networks using ZeroCostDL4Mic and (ii) performing inference with deepImageJ, following the export of these models to the bioimage.io format. Within the deepImageJ environment, an ImageJ macro is utilized to process the five available time points of Lifeact-RFP images with Pix2Pix. This step results in synthetic SiR-DNA images, which effectively stain the nuclei. Subsequently, these images undergo processing with the deepImageJ implementation of StarDist, which involves applying the UNet model (trained via ZeroCostDL4Mic) in Fiji/ImageJ and the corresponding post-processing for nuclei segmentation to produce masks. These masks, obtained from five distinct time points, are then tracked using TrackMate^(18)^, an ImageJ plugin, to visualize cell trajectories and track cell movement.

**Figure 1.**
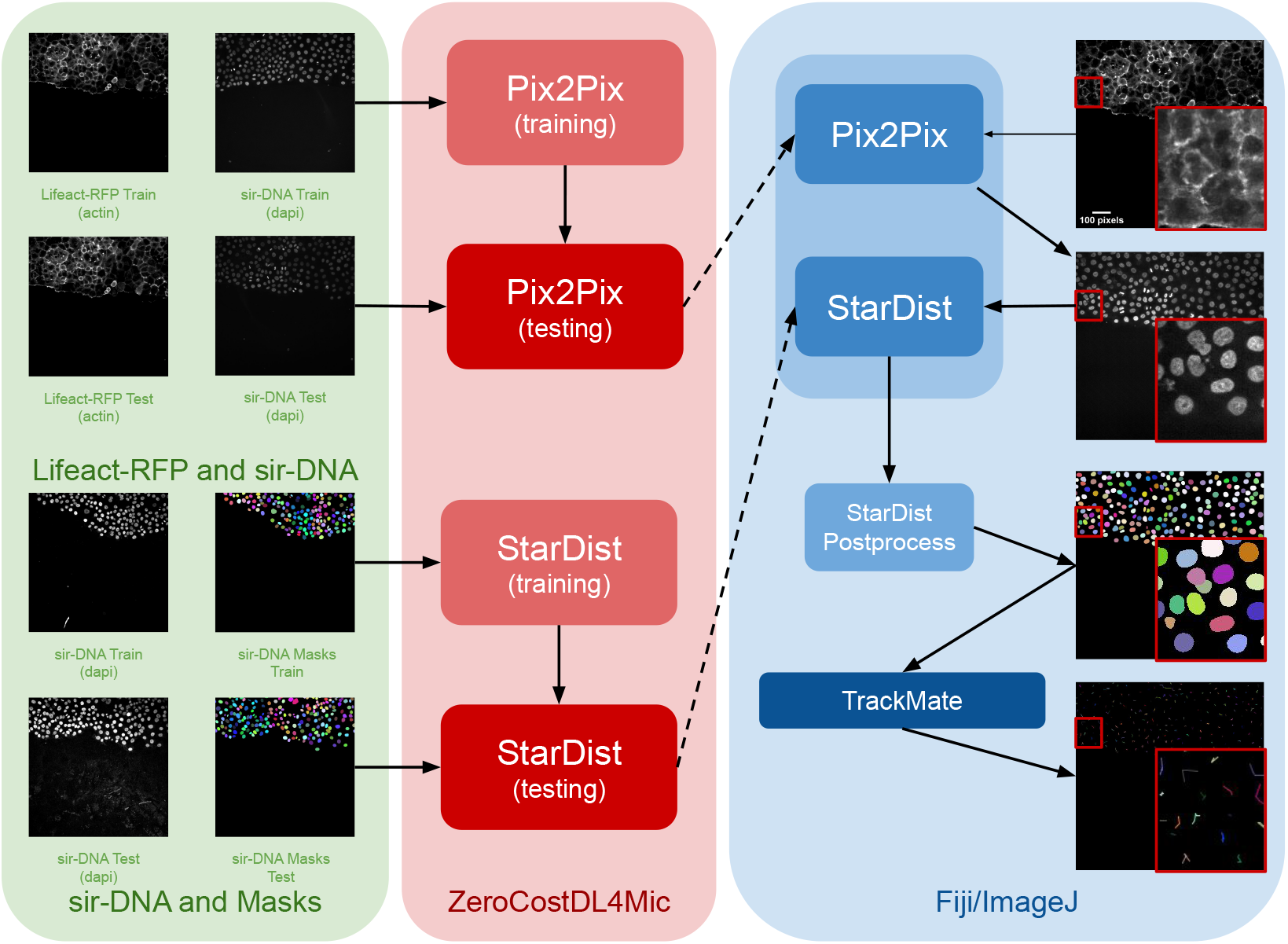
Case Study 1: Image-to-Image translation and cell Segmentation Pipeline and Dataset. The pipeline involves three main stages: dataset preparation, model training using ZeroCostDL4Mic, and inference and post-processing in deepImage. Initially, Pix2Pix and StarDist are fine-tuned with specific datasets. Pix2Pix transforms actin images into synthetic DAPI images, while StarDist creates masks from DAPI images. Once trained, the models are exported to the BioImage Model Zoo format and subsequently installed in deepImageJ. In the Fiji/ImageJ and deepImageJ environment, the pipeline first uses Pix2Pix to transform actin images into synthetic DAPI images, followed by the application of StarDist for nuclei segmentation. Finally, TrackMate is utilized for a thorough evaluation of cell tracking. A contrast enhancement has been applied to actin images for visualization purposes..

### 3.2. Case Study 2: Comprehensive 3D Nuclei Segmentation with deepImageJ

Case Study 2 emphasizes the capabilities of deepImageJ, particularly benefiting from its integration within the extensive image processing ecosystem of Fiji/ImageJ. This integration affords the flexibility to run advanced pipelines automatically, including 3D+t image analysis, in a user-friendly manner. In particular, Case Study 2 demonstrates the segmentation of nuclei in microscopy images of whole embryos.

For enhanced reproducibility and to accommodate users without access to high-powered computational resources, this pipeline is executed in 2D, using a lightweight framework. By employing StarDist 2D (UNet model + postprocessing) and then applying MorphoLibJ^(19)^ connected components in 3D, we successfully mimic 3D segmentation. This approach demonstrates how the integration of deepImageJ into the Fiji/ImageJ ecosystem facilitates complex image analysis tasks, bypassing the need for extensive computational power typically required for direct 3D processing in bioimage analysis.

The dataset for this study is part of the Cell Tracking Challenge repository, specifically the “Developing *Tribolium Castaneum* embryo”^(20)^. This dataset provides 3D volumetric data of two beetle embryos, complete with accompanying sparse nuclei annotations of the beetle’s blastoderm at the junction of embryonic and extra-embryonic tissues. Several preprocessing steps are undertaken to leverage deepImageJ’s capabilities for running StarDist 2D. Initially, a targeted selection of slices from various timepoints in embryo 01 was conducted, guided by the availability of ground truth data within the dataset from the Cell Tracking Challenge. This selection process was governed by the necessity to choose slices for which ground truth data existed, as these were imperative for the training phase. These images and masks are then downsampled to manage memory usage and ensure reproducibility. Following this, a median filter (kernel size, 7 pixels) is applied to reduce noise in the input images. The prepared pair of images is then processed using the StarDist notebook within the ZeroCostDL4Mic repository, as illustrated in Figure 2.

**Figure 2.**
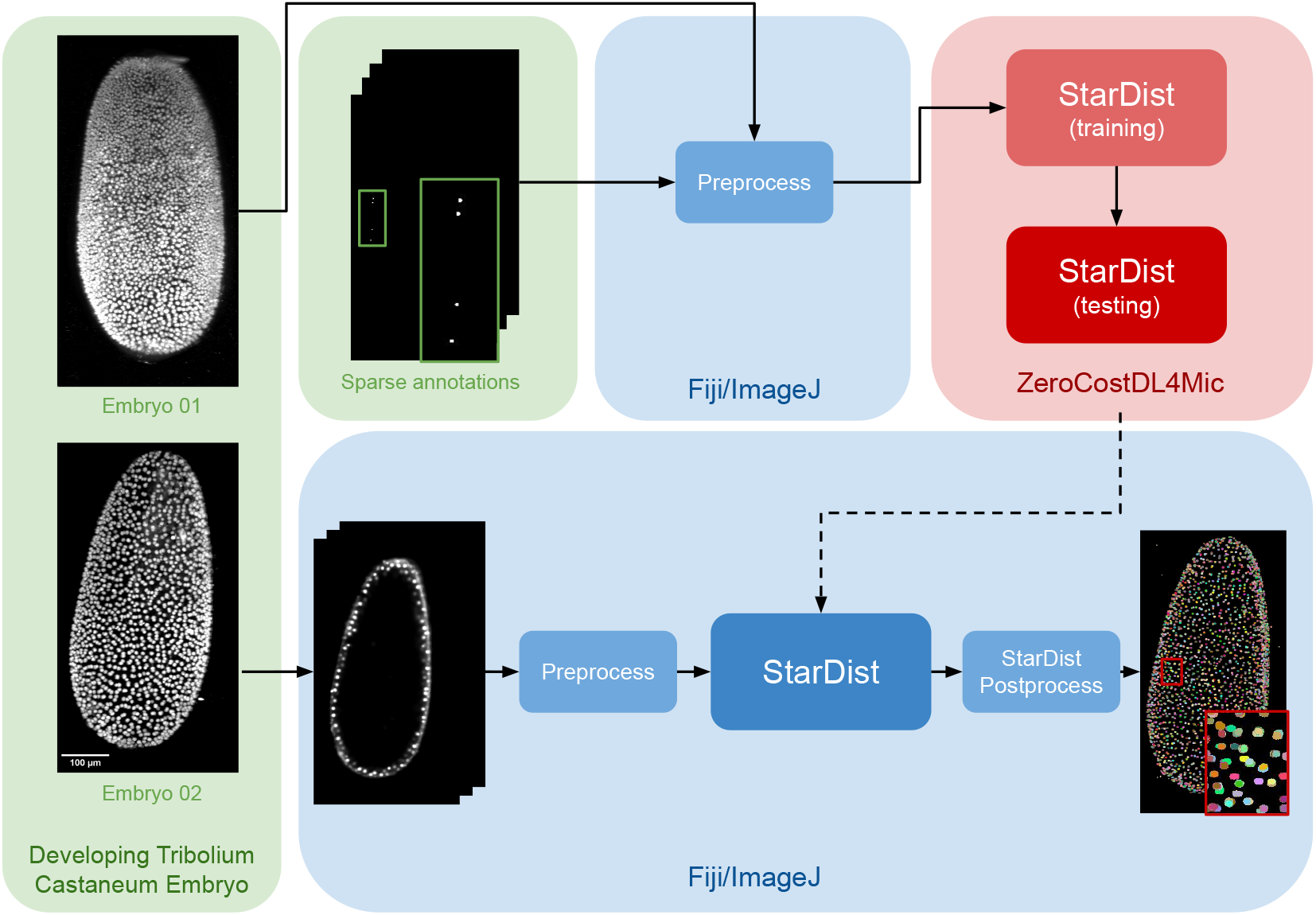
Case Study 2: 3D Nuclei Segmentation Pipeline and Dataset. The dataset consists of two distinct embryos, labeled 01 and 02. One embryo is used for fine-tuning the StarDist network in ZeroCostDL4Mic, following downsampling and noise filtering, while the other is utilized for inference. After training the StarDist model, it is employed in deepImageJ to create the masks, followed by StarDist postprocessing. The pipeline is completed with the application of Connected Components for 3D visualization. All 3D volumes are displayed as Z-projections..

In Case Study 2, after the UNet model is fine-tuned, it is exported and integrated into deepIm-ageJ as a bioimage.io model. The subsequent analysis in Fiji/ImageJ involves a structured approach: i) implementing preprocessing steps that mirror those used during the model’s training, ii) deploying the StarDist model for inference, and iii) applying a series of post-processing techniques for assessment and visualization. This includes downsampling and denoising of selected timepoints from Embryo 2, following the methodology utilized for the training dataset (Embryo 01). Each 2D slice of the embryo is then processed through the trained network. The final step involves enhancing the segmentation masks using the StarDist post-processing pipeline and applying MorpholibJ’s Connected Components^(19)^ for comprehensive 3D visualization of the nuclei.

### 3.3. Case Study 3: Segmentation of Arabidopsis Apical Stem Cells and Integration with the BioImage Model Zoo in deepImageJ

In this use case, we highlight two key capabilities of deepImageJ: its adeptness in handling large 3D images and its seamless integration with the BioImage Model Zoo.

The implementation of this pipeline involves using the 3D UNet Arabidopsis Apical Stem Cells model from the bioimage.io website^(21)^, paired with the dataset titled “Research data supporting Cell size and growth regulation in the Arabidopsis thaliana apical stem cell niche”^(22)^. This approach establishes an efficient yet robust pipeline for cell segmentation within apical stem cells, particularly focusing on the epidermal cell volumes in the apical meristem, using the 3D UNet pretrained model from the BioImage Model Zoo.

The main steps of the pipeline are summarized in Figure 3. Initially, the model is downloaded from bioimage.io website and installed via the deepImageJ Install mode. Subsequently, a relatively large image, measuring 515 × 515 pixels with 396 slices and representing a 3D volume of plant 13 (chosen for its significant size), is selected for the analysis. The 3D UNet is then employed, and using the tiling strategy of deepImageJ^(12)^, the image is processed in 180 patches of 100 × 128 × 128 pixels to cover the whole 3D volume. After running the model and obtaining the mask, the Gamma function at 0.80 is applied to enhance the membranes. Then, The morphological segmentation GUI ofMorphoLibJ^(19)^ is executed. Namely, a combination of morphological operations (extended minima and morphological gradient) previous to a watershed flooding algorithm is applied with a low tolerance setting to guarantee precise segmentation accuracy.

**Figure 3.**
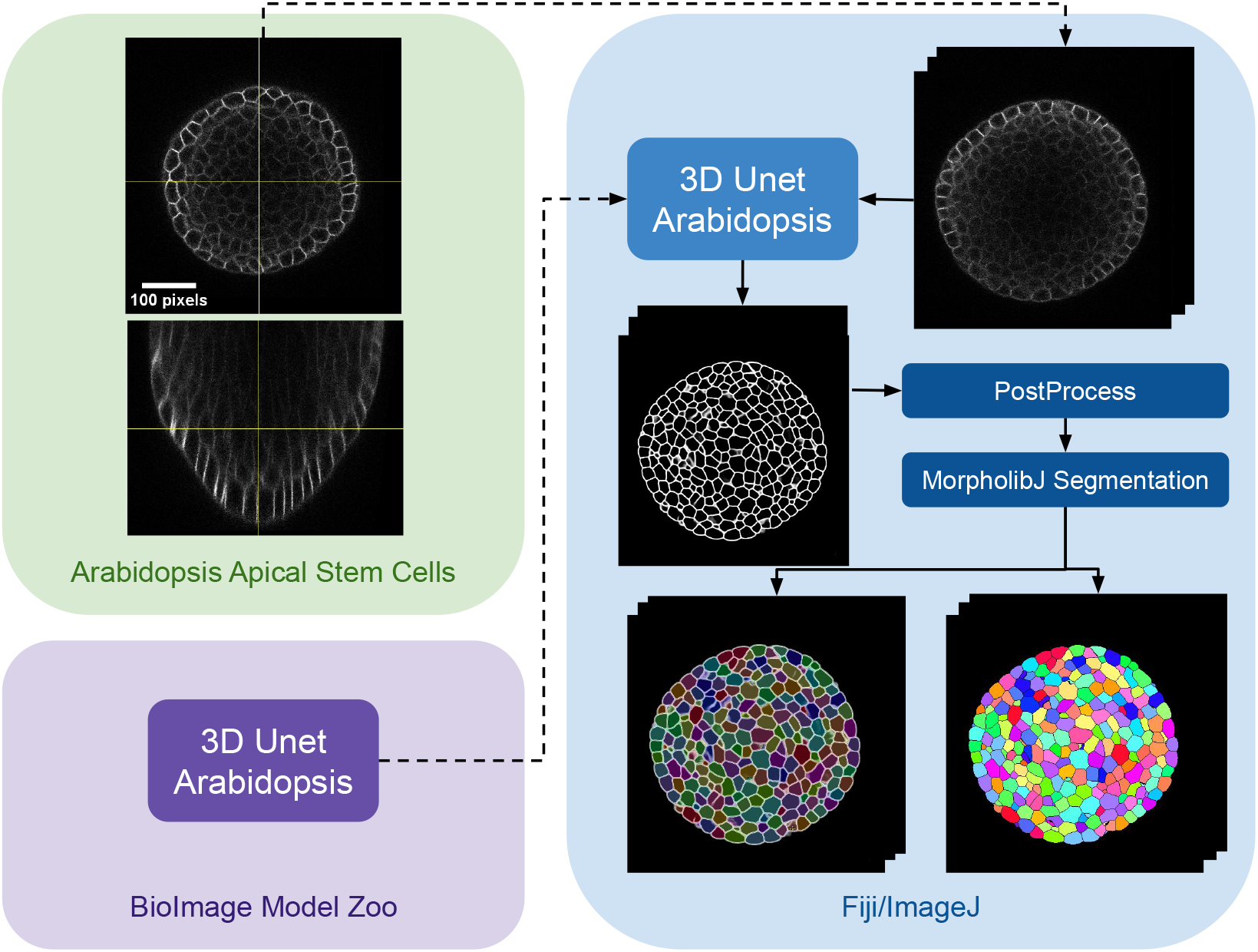
Case Study 3: Segmentation of Arabidopsis Apical Stem Cells - Pipeline.This diagram illustrates the pipeline for Case Study 3. Initially, the dataset is acquired, followed by downloading and installing the model from the BioImage Model Zoo into deepImageJ. Subsequently, the model is applied to a selected root volume to generate a mask. The process concludes with post-processing and MorpholibJ segmentation to display catchment and overlay basins on the segmented image..

## 4. Discussion

The advancements presented in this manuscript reflect a significant leap in the capabilities of deepImageJ, a key plugin for Fiji/ImageJ in the domain of bioimage analysis. This tool evolution has the potential to positively impact the life sciences community, where the need for accessible, efficient, and versatile image analysis tools is ever-growing. Integrating deepImageJ with the BioImage Model Zoo and incorporating the JDLL underscore its role as a bridge between complex DL models and practical, user-friendly applications. Moreover, the seamless integration with the JDLL enhances the software’s capabilities, providing a unified platform for deploying diverse DL models.

The presented case studies demonstrate the profound adaptability and enhanced functionality of deepImageJ. A summary of these case studies is depicted in Figure 4. The first case study, focusing on image-to-image translation and nuclei segmentation, illustrates the software’s ability to integrate and execute multiple DL environments within a single Fiji/ImageJ instance. This capability is crucial in biological contexts where multi-faceted analysis is often required. It facilitates the researchers to delve into intricate biological pipelines without the need for extensive coding expertise. The second case study further showcases the power of the integration of deepImageJ into the Fiji/ImageJ ecosystem to, in this case, handle complex 3D nuclei segmentation. A task that is increasingly relevant as imaging technologies advance. Finally, the third case study emphasizes the tool’s adeptness in processing large 3D images, an essential feature for analyzing extensive datasets commonly encountered in modern biological research as well as the deepImageJ integration with the BioImage Model Zoo.

**Figure 4.**
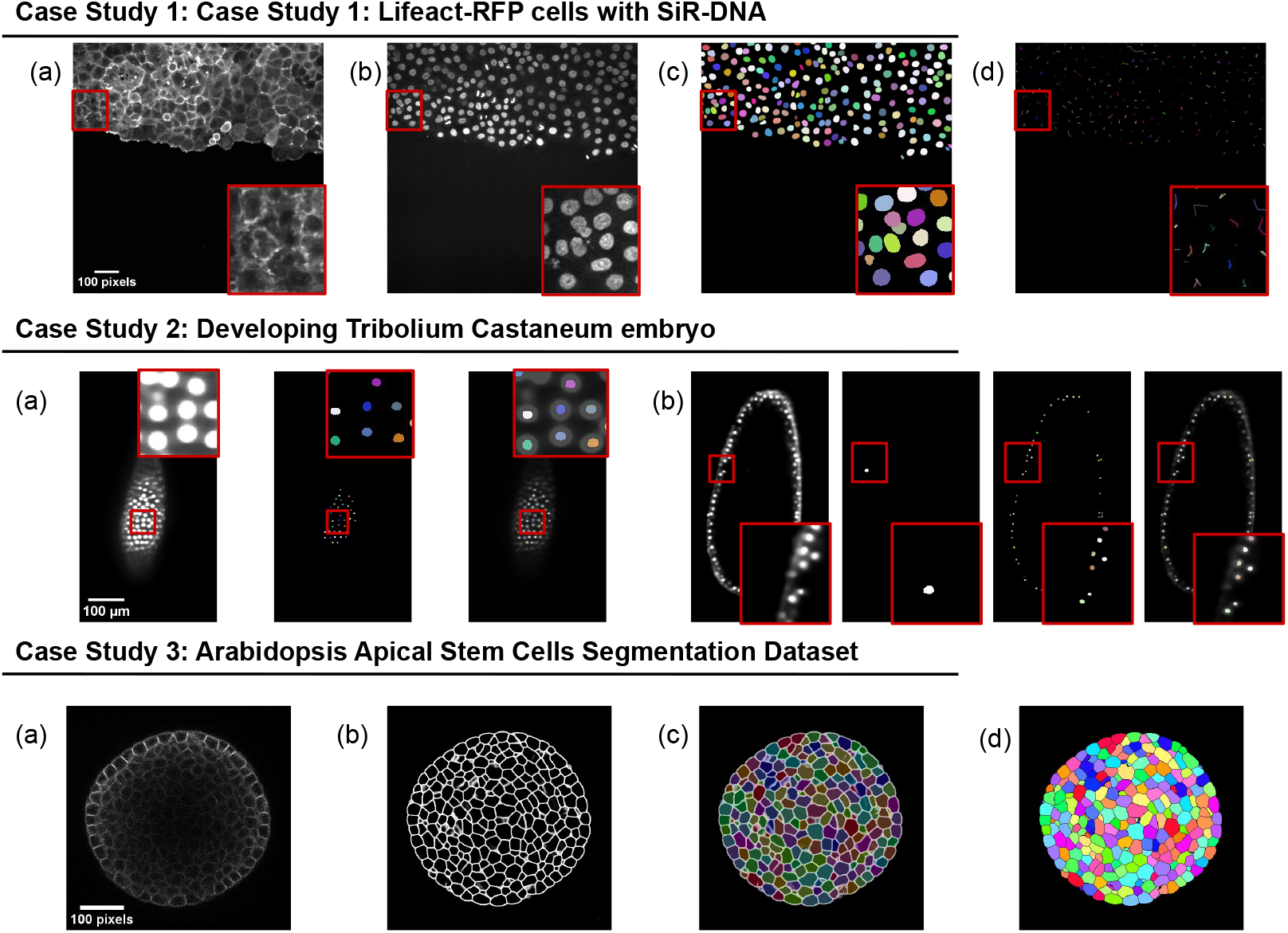
Summary for the three case studies. This figure provides an overview of three distinct case studies, highlighting deepImageJ’s versatility and integration with other tools and plugins. Case Study 1 illustrates the transformation of an actin membrane stain image into a synthetic nuclei stain image via Pix2Pix, followed by StarDist nuclei segmentation and TrackMate cell tracking. Case Study 2 presents two examples of a single slice from input volume and StarDist output, with one including the Ground Truth. Case Study 3 shows the pipeline stages: (a) input image, (b) mask generation, (c) overlay of Morphological Segmentation basins, and (d) visualization of catchment basins..

Looking ahead, a key area of focus for future work is the integration of interactive annotation features. This functionality will enable users to fine-tune DL models directly within the deepImageJ environment. By incorporating tools for interactive annotations, researchers will have the flexibility to customize and refine their models with greater precision and ease, tailoring them to specific research needs. This addition is particularly crucial for cases where standard pre-trained models may not perfectly align with unique dataset characteristics, allowing for more personalized and accurate analysis.

Another significant advancement planned is the implementation of a transparent connection with Python. This development will endow deepImageJ with full training capabilities, effectively transforming it into a comprehensive platform for both model development and application. By bridging deepImageJ with Python, a leading language in the field of data science and machine learning, users will gain access to a vast ecosystem of libraries and tools. This integration will not only facilitate the training of DL models within deepImageJ but also enable seamless interoperability between deepImageJ and a wide range of Python-based data processing and analysis frameworks.

The development of the deepImageJ environment and related initiatives represent a paradigm shift in how biologists and researchers approach bioimage analysis. By lowering the barrier to entry for applying advanced DL techniques, deepImageJ democratizes access to cutting-edge analysis methods. This accessibility is vital for fostering a culture of innovation and exploration in the life sciences, where researchers can leverage these tools to uncover new insights into complex biological phenomena. As bioimage analysis continues to evolve, tools like deepImageJ will play a crucial role in bridging the gap between advanced computational techniques and practical research applications, driving forward the frontiers of science and medicine.

## 5. Conclusions

In conclusion, this manuscript has presented the evolution of deepImageJ, highlighting key advancements and new features. The integration of the JDLL has played a pivotal role in expanding the capabilities of deepImageJ, making it a versatile and accessible tool for life scientists and bioimage analysts. The case studies showcased the practical applications of deepImageJ across different biological scenarios, demonstrating its effectiveness in tasks ranging from cell segmentation to plant tissue analysis.

The introduction of the JDLL has significantly streamlined the execution of DL models, providing a unified framework for various DL engines and frameworks. The ability to run different engines in a single Fiji/ImageJ instance opens up new possibilities for constructing complex image analysis pipelines. The enhanced compatibility with TensorFlow 1 and 2, PyTorch, and ONNX, coupled with the capability to process larger images, marks a significant step forward in the field of bioimage analysis.

The zero-coded nature of deepImageJ, coupled with the new features introduced in version 3.0, underscores its commitment to democratizing access to DL tools for life scientists. This paper not only serves as a comprehensive documentation of deepImageJ’s journey but also aims to inspire the research community to harness the power of DL in the realm of bioimage analysis.

## 6. Materials

### 6.1. Datasets

#### 6.1.1. Case Study 1: Lifeact-RFP cells with SiR-DNA

The datasets employed for training in both tasks are publicly available and have been used extensively for similar research in fluorescence microscopy. These datasets consist of images of live cells expressing Lifeact-RFP (Red Fusion Protein) for visualizing actin filaments and are treated with 0.5 µM SiR-DNA for live cell DNA staining. The continuous imaging of cell culture was performed over 14 hours using a spinning disk confocal microscope, capturing images at 10-minute intervals. This imaging was done with a Yokogawa CSU-W1 scanning unit on an inverted Zeiss Axio Observer Z1 microscope with a 20x (NA 0.8) air, Plan-Apochromat objective (Zeiss)^(9,23)^.

For the Pix2Pix network training, we used pairs of images from these datasets: membrane-stained (Lifeact-RFP) and nuclei-stained (SiR-DNA) images. The same nuclei-stained images were also employed for training the StarDist model^(23)^. Manual mask images were generated in Fiji/ImageJ^(9)^ to outline each nucleus, using the freehand selection tool to trace and add outlines to the Region of Interest (ROI) manager, followed by the creation of an ROI map with the LOCI plugin. These ROI maps were crucial for the accurate training of the StarDist model, particularly following image-to-image translation.

#### 6.1.2. Case Study 2: Developing Tribolium Castaneum embryo

In the context of Case Study 2, a specialized dataset from the Cell Tracking Challenge was utilized to fine-tune the StarDist network (see Supplementary Material for the link to donwload). This dataset includes high-resolution fluorescence microscopy images capturing the developing *Tribolium Castaneum* embryo nuclei. The images were acquired using a Zeiss LightSheet LZ.1 microscope equipped with a Plan-Apochromat 20x/1.0 (water) objective lens, achieving a voxel size of 0.38 × 0.38 × 0.38 microns. The images were taken at 1.5-minute intervals to track cellular dynamics over time. For detailed information on sample preparation, RNA injections, and imaging techniques, please refer to Jain etal. ^(24)^.

It is important to recognize that the image data, acquired using a light-sheet microscope, was fused from multiple viewpoints. Due to this, some views might not align perfectly, leading to the appearance of false or conspicuous nuclei. Furthermore, as not all views cover the entire volume, localized dark patches may be present along the image axis.

The experiment involved two separate embryos. The first embryo (Embryo 01) was used to train the StarDist network in ZeroCostDL4Mic, whereas the second (Embryo 02) served as an independent dataset for testing the methodology in Fiji/ImageJ. Crucially, the annotations used for network training are sparse, focusing only on selected regions and cell lineages. This sparsity, particularly in the beetle’s blastoderm at the junction of embryonic and extra-embryonic tissues, was pivotal for effective network training. The sparse annotations provided a focused and relevant dataset for fine-tuning the network, as depicted in Figure 2.

#### 6.1.3. Case Study 3: Arabidopsis Apical Stem Cells Segmentation Dataset

In this case study, we utilized a publicly available confocal imaging-based dataset of plant cells from Willis et al. ^(22)^, which includes data from six Arabidopsis thaliana plants treated with Naphthylphthalamic acid (NPA). This treatment inhibits auxin transport, allowing the study of its effects on plant development and physiology.

Confocal z-stacks were acquired every 4 hours for 3-3.5 days at a resolution of 0.22×0.22×0.26*μm*^3^ per voxel using a 63 × 1.0 N.A. water immersion objective. For each plant, approximately 20 data time points were available. Each time point comprises a stack of around 200 image slices, with each slice measuring 512 × 512 pixels. The dataset includes segmentation ground truth, for instance, segmentation of each cell. Specifically, we analyzed an image stack from plant number 13, which displays cell membranes expressing acylYFP in a shoot apical meristem 84 hours post-treatment.

## 7. Methods

### 7.1. Case Study 1: Lifeact-RFP cells with SiR-DNA

#### 7.1.1. Pix2Pix for Image Translation and StarDist for nuclei segmentation

The Pix2Pix model^(16)^, integral to Case Study 1 for the task of image-to-image translation from membrane staining (Lifeact-RFP) to nuclei staining (SiR-DNA), underwent rigorous training for 200 epochs.

The training dataset consisted of 1748 paired image patches, each with dimensions (1024 × 1024 × 3), and a patch size of (512 × 512). The training process utilized a batch size of 1 and a vanilla Generative Adversarial Network (GAN) loss function. Executed within the Pix2Pix ZeroCostDL4Mic notebook (v1.15.1) on Google Colab, the model was trained to adhere to the default parameters of the notebook. No data augmentation was applied during training. Key training parameters encompassed a patch size of 512 × 512, a batch size of 1, and an initial learning rate of 2*e* − 4, achieving successful translation from membrane to nuclei staining. The Pix2Pix model is exported using PyTorch 2.0.1.

After the image-to-image translation, the StarDist model, designed for nuclei segmentation, underwent extensive training for 100 epochs. StarDist consists of a UNet trained to identify the intrinsic features of an object, such as the centroid or oriented distances from the centroid to its boundary, that enable its reconstruction as a star-convex polygon. The 2D variant of StarDist^(11)^ was trained and evaluated using its implementation within the StarDist 2D ZeroCostDL4Mic notebook (v 1.19). The training dataset consisted of 45 paired image patches, each with dimensions (1024 × 1024), and a patch size of (1024 × 1024). The training process used a batch size of 2. The model was fine-tuned from a pretrained model, applying no data augmentation during training. Executed within the Google Colab environment, the training parameters included a patch size of 1024*x*1024, a batch size of 4, 100 epochs, and an initial learning rate of 3*e* − 4. The resulting StarDist model is exported with Tensorflow 2.14.

#### 7.1.2. Post-Processing with StarDist and TrackMate

In Case Study 1, the reconstruction of 2D star-convex polygons is facilitated by the StarDist plugin for Fiji/ImageJ, which supports macro recording in ImageJ. Therefore, an ImageJ/Fiji macro is utilized to execute the complete pipeline, encompassing the running of Pix2Pix, StarDist, and the subsequent StarDist PostProcessing. Following the generation of masks by this pipeline, the trajectories of cells across five available timepoints are analyzed using TrackMate^(18)^. The results, including the cell trajectories, are illustrated in Figure 1.

### 7.2. Case Study 2: Developing Tribolium Castaneum embryo

#### 7.2.1. Preprocessing

In Case Study 2, preprocessing steps are essential before inputting images into the StarDist network. It is important to note that for fine-tuning the ZeroCostDL4Mic model, we utilize sparse annotations of the beetle’s blastoderm, as described in 6.1.2. Consequently, only selected slides from the dataset are employed. During the inference process in deepImageJ, however, the entire volume corresponding to each time point is processed.

A two-stage preprocessing strategy is implemented to address the image noise and reduce the computational load. The first step involves applying a median filter across all images to reduce noise effectively. Following this, a downsampling operation is conducted. This operation reduces the resolution by half along the *x* and *y* axes for the slices used in fine-tuning the ZeroCostDL4Mic model and along all three axes (*x, y*, and *z*) when processing the entire volume for inference with deepImageJ.

#### 7.2.2. StarDist - Nuclei Segmentation in 2D

The segmentation network employed in Case Study 2 is based on StarDist^(11)^, a deep-learning method designed to precisely segment cell nuclei from bioimages. This method uses a shape representation founded on star-convex polygons to predict both the presence and shape of nuclei within an image. The 2D variant of StarDist relies on an adapted UNet architecture, allowing for efficient segmentation of 2D datasets. Implemented within the ZeroCostDL4Mic framework, the StarDist 2D model was specifically tailored for nuclei segmentation within the context of the Developing *Tribolium Castaneum* embryo dataset, as described in Section 6.1.2. The dataset structure was adjusted accordingly to facilitate compatibility with the notebook’s data reading mechanism. The code detailing the data structuring process is available on the deepImageJ GitHub for reproducibility purposes. Dataset augmentation was performed by a factor of four via random rotations, flips, and intensity changes.

The training regimen involved 50 epochs on 40 paired image patches of size 512 × 512 cropped from the original images (1871×965 pixels). A batch size of 15 was utilized, employing a MAE loss function. The model was retrained from a preexisting pretrained model (2D Versatile fluo from StarDist Fiji), with key training parameters including a learning rate of 5*e* − 05, 10% validation data, 32 rays (n_rays), and a grid parameter of 2. Despite challenges associated with ground truth variability, particularly in cases where only one nucleus is marked in the mask, the model demonstrated good performance. This effectiveness was observed in the segmentation accuracy, where the predicted results were consistently aligned with the available ground truth, despite its inherent variability.

#### 7.2.3. Postprocess with StarDist

In Case Study 2, the post-processing of StarDist is streamlined through an ImageJ macro, which processes each slice of the 3D embryo volume independently. The macro, designed to handle 488 slices per timepoint, applies the StarDist model slice by slice. For each slice, the StarDist model is applied, followed by a series of post-processing operations. These operations, crucial for accurate object detection and minimizing overlap, include applying specific thresholds for probability and non-maximum suppression. The macro utilizes the StarDist plugin to ensure precise segmentation results for each 2D slice.

This macro effectively transforms the multi-channel output of StarDist into a single, comprehensive mask. In the final phase, the Connected Components algorithm is applied across the entire 3D volume. This process results in a detailed visualization of the entire volume, with each segmented cell clearly delineated, as illustrated in Figure 2.

### 7.3. Case Study 3: Arabidopsis Apical Stem Cells Segmentation Dataset

#### 7.3.1. 3D UNet Arabidopsis Apical Stem Cells

In Case Study 3, a 3D UNet^(21)^ was employed for cell boundary segmentation. This pre-trained model is accessed from the bioimage.io website for inference, and it is also available through zenodo^1^.

The authors of the network^(21)^ employed a training strategy where the 3D UNet was trained on ground truth cell contours obtained by applying a Gaussian blur to a two-voxel-thick boundary between labeled regions. The training regimen featured a combination of binary cross-entropy (BCE) and Dice loss, with notable architectural modifications, including replacing batch normalization with group normalization and utilizing same convolutions instead of valid convolutions. During training, augmentation techniques such as flips, rotations, elastic deformations, and noise augmentations were employed to enhance the model’s generalization capabilities.

The trained model is available on bioimage.io under the `emotional-cricket`, allowing accessibility for the wider research community. In this case study, the network is exclusively employed for inference on the specified dataset.

#### 7.3.2. PostProcess

In the post-processing phase of Case Study 3, the pipeline includes two distinct steps. Initially, a Gamma correction function, set at 0.80, is applied to enhance membrane visibility and reduce any blurriness. Subsequently, the Morphological Segmentation tool from MorpholibJ^(19)^ is utilized for both segmentation and visualization. This tool is employed with a tolerance setting of 10, enabling the effective depiction of catchment and overlay basins on the segmented image. This precise application of Morphological Segmentation ensures clear and distinct visualization of each cell.

## Acknowledgements

We thank Lucia Moya-Sans and Ivan Estevez for their effort in the BioImage Model Zoo activities related to deepImageJ. The authors acknowledge the assistance of ChatGPT, provided by OpenAI, for its role in reviewing and correcting grammatical errors within this manuscript, ensuring clarity and coherence in the presentation of the research findings.

## Accessibility of Data, Code, and Models

To ensure the reproducibility of our study, we have made all data, code, and models accessible in various formats. The datasets are accessible from the dedicated repositories specified in the Datasets section. For models fine-tuned during the training process, they are accessible in the BioImage Model Zoo and Zenodo, and the corresponding notebooks used for fine-tuning are available on ZeroCostDL4Mic. Additionally, all codes used for dataset construction, ImageJ Macros, and other relevant tools are accessible in a GitHub repository associated with deepImageJ. All links and details are also specified in the Supplementary Material as well as in the GitHub repository dedicated to it at https://github.com/deepimagej/case-studies.

## Ethical Standards

This study adheres to ethical principles and guidelines in scientific research. We utilized publicly available datasets in compliance with their terms and conducted no live experiments, thus negating the need for institutional ethical approval. The study upholds standards of integrity, transparency, and reproducibility, ensuring accurate and responsible reporting. No human participants, animal experiments, or identifiable personal data were involved in this research.

## Author Contributions

All the authors contribute to the conception of the project. C.F.B, C.G.L.H and E.G.M wrote the software code with input from W.O.; C.F.B performed the experiments with input from E.G.M, D.S, and A.M.B; E.G.M designed the pipelines with input from D.S and A.M.B; D.S and A.M.B supervised the research; C.F.B and A.M.B wrote the manuscript. All the authors read and approved the final submitted draft.

## Supplementary Material

State whether any supplementary material intended for publication has been provided with the submission.

## Supplementary Material: Data, Notebooks, Code, and Models Availability

The following table provides details on the datasets, notebooks, code, and models used in each case study. This information is essential for ensuring reproducibility and is included in the supplementary material.

**Table 1.** Availability of data, notebooks, code, and models for each case study.

| Case Study | Resource Type | Description and Links |
| --- | --- | --- |
| Case Study 1 | Files | prepare_dataset.py, StarDist Postprocess macro CS1.ijm |
|  | Notebooks | Pix2Pix Notebook, StarDist 2D Notebook |
|  | Dataset | Lifeact-RFP, sir-DNA DAPI |
|  | Models | Pix2Pix Model, StarDist Model |
| Case Study 2 | Files | Generated_GT.py, Mount_stardist_dataset.py, StarDist_postprocess_macro_cs2.ijm |
|  | Notebooks | StarDist 2D Notebook |
|  | Dataset | Developing Tribolium Castaneum Embryo |
|  | Model | StarDist Model |
| Case Study 3 | Dataset | Arabidopsis Apical Stem Cells |
|  | Model | 3D Unet Arabidopsis Model |

Additionally, all codes related to these case studies can be found in the GitHub repository deepImageJ Case Studies.

## Footnotes

1. 3D Unet Arabidopsis Apical Stem Cells Model in Zenodo https://zenodo.org/records/7768142

